# Modeling of Internal and External Factors affecting a Complex Dengue Network

**DOI:** 10.1101/2020.12.01.406033

**Authors:** Hafiz Abid Mahmood Malik, Faiza Abid, Mohamed Ridza Wahiddin, Ahmad Waqas

## Abstract

There are different factors that are the cause of abrupt spread of arbovirus. We modelled the factors (internal & external) that can increase the diffusion of dengue virus and observed their effects. These factors have influenced on the *Aedes aegypti* (a dengue virus carrier); factors which increase the dengue transmission. Interestingly, there are some factors that can suppress the *Aedes aegypti*. The species of *Aedes aegypti* formalizes its own network by which dengue virus is spread. Internal & external exposures of the dengue epidemic complex network have been modelled and analyzed. Influence of internal and external diffusion with two scenarios has been discussed. ‘Genetically modified mosquito’ technique has been applied and its associated simulated results are discussed. From the outcomes, the best time duration to contain the spread of dengue virus has been proposed, and our simulation model showed the possibility of suppressing the *Aedes aegypti* network.

## 1. Introduction

Because of the danger posed by the *Aedes aegypti*, there have been many attempts to control its populations. Unfortunately, in spite of the best endeavors of governments and groups, these endeavors have, up to this point, not been exceptionally fruitful and there are presently no authorized antibodies or particular therapeutics to control this mosquito and its virus [1] [2] [3] [4].It is important to mention here that *Aedes aegypti* is also the primary vector/carrier of chikungunya, yellow fever and *Zika* virus (ZIKV). ZIKV has become a serious threat in the tropical zones in the world. Furthermore, CoViD-19 is very critical for its transmission channels. So, this study is also beneficial for CoViD-19, ZIKV and other diseases that has the spreading transmissions like person-to-person. The topology of biological networks is an important factor to be considered. [5]

Female *Aedes aegypti* is the dengue vector, meaning that only the female mosquito of this species is able to spread the virus [6] [7]. A female *Aedes aegypti* mates with a male mosquito in order to grow its generation through laying eggs (a natural way). It is considered one of the internal factors to increase the dengue network. However, its virus might be transferred by the infected persons (Fig.1), if any other mosquito (not the species of *Aedes aegypti*) bites the infected human and may become a source of dengue virus [8] [9].It is also another source of external factor to increase the dengue complex network. Therefore, there is a need to break down this chain network by inserting some exogenous factors.

**Fig. 1.**
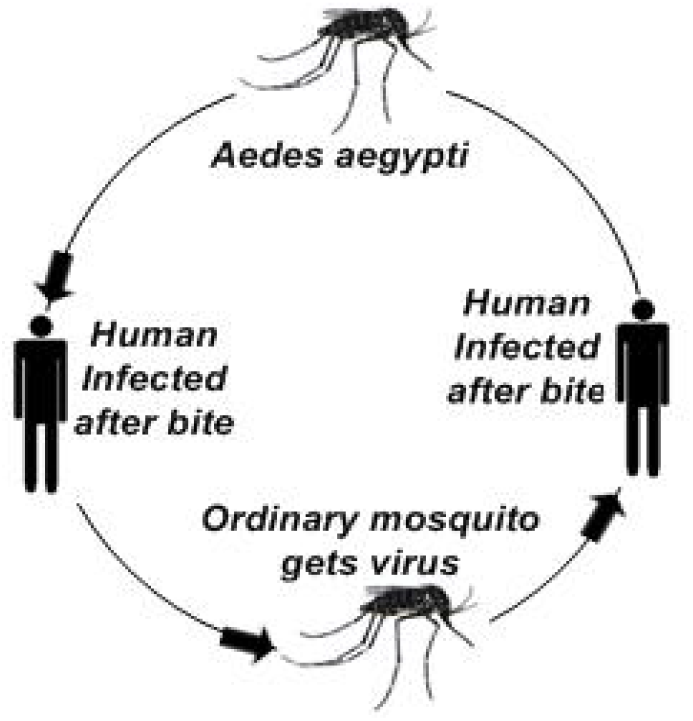
Transmission of Dengue virus.

In this study, a dataset of the Dengue Virus (DENV) affected cases from the Ministry of Health (MOH) Malaysia has been used [10] [11]. The dataset of Selangor (a state of Malaysia) has been obtained, in which a weekly number of dengue cases are recorded in all affected localities from the periods of 20 October 2013 to 18 October 2014. Here, weekly number of dengue cases are edges and affected localities are nodes, that constitute the dengue epidemic network of interest. Furthermore, there are 560 affected nodes with 36,878 dengue cases [10]. Dataset (*Appendix B*) has been modelled into two-mode network and then projected in one-mode network, and after applying different metrics, results have been used in this study [12] [13][14] [15]Moreover, in this paper we modelled the internal and external factors and observed their influence on the transmission of dengue virus. It is considered that *Aedes aegypti* has geographically organized itself in Selangor Malaysia to form a network. Here, the ‘network’ is the linkages of *Aedes aegypti* with other factors to transmit its infections. There are a variety of approaches ranging from sterile insect technique (SIT) to Wolbachia which prevents transmission. This ‘genetically modified mosquito’ technique, has been applied in this research. It produced (in simulated results) more efficient outcomes in the scale-free network through targeted attacks in the focal hubs of *Aedes aegypti*. It has been proven that dengue epidemic follows a scale-free network [16] [17]. The outcomes from this study are important for researchers, health officials and decision makers who deal with the arbovirus epidemics, like *Zika* virus, chikungunya, and yellow fever. There are various examples for complex network [18] that gave motivation to model the dengue complex network phenomenon. For CoViD-19 (Corona Virus Disease), transmission is increasing due to person-to-person close contact, so internal and external factors need to be controlled.

## 2. Research Methodology

We divide the methodology into two aspects; the possible causes of DENV spread and any possible treatments. This is considered as: *Scenario 1* and *Scenario 2*.

### Scenario 1

What factors can be a cause of the diffusion of dengue virus? For example; untidy spots, water-holding containers, used tires containing rain water, pots of plants and flower beds. All of these are standard reasons. Within the dengue epidemic network two cases are discussed in this paper: influence of internal exposure and influence of external exposure. In first case, influence of internal exposure means the source of dengue virus diffusion is present in the same cluster of complex networks. In Fig.2, let’s consider Gombak (a district of Selangor, Malaysia) is a cluster (*Ci*) of *i* nodes. Where *Ci* contains some infected nodes (such as: *Li2, Li3, Li4, Li7*) and uninfected nodes (such as: *Li1, Li5, Li6, Li9*). Suppose, an *Aedes aegypti* bites a person in the node *Li3*, and now that person may be a cause of DENV spread to the next-door neighbours within *Li3* or in the other node such as *Li5* (see Fig.2). Similarly, *Li7* is an infected node in the *Ci* that may infect the *Li6* which remains to be uninfected. It is referred as internal factor diffusion and called case 1. As well as this, if any *Aedes aegypti* spreads its infections to more than one person within the same cluster, it is also referred to as case 1.

**Fig. 2.**
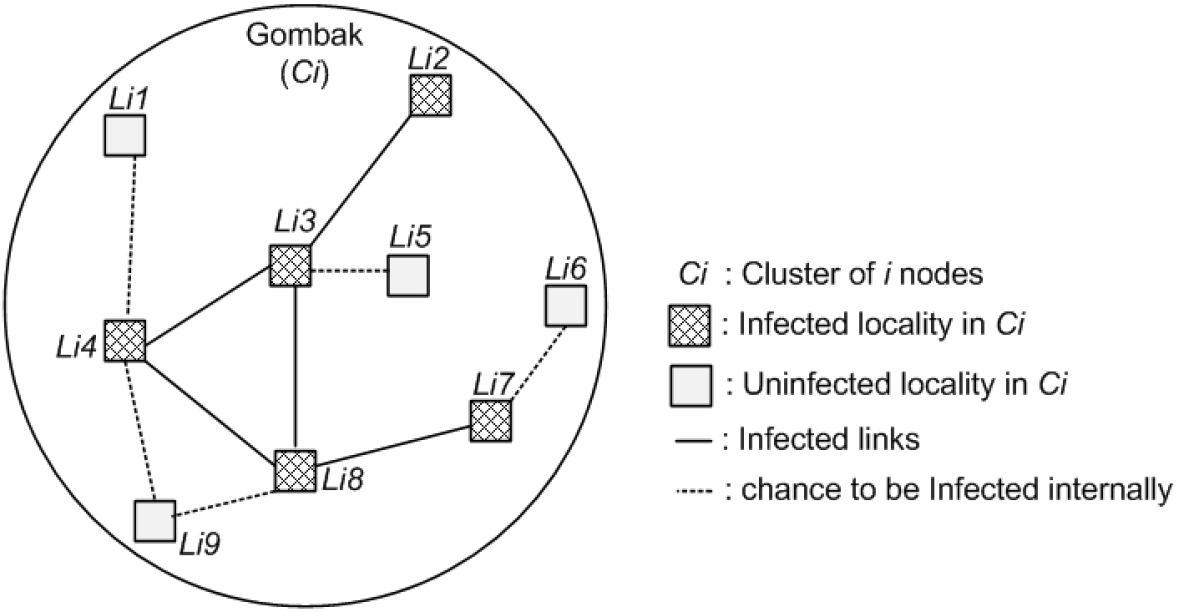
Influence of Internal diffusion (Scenario 1: Case 1), based on the given dataset.

In second case, influence of external exposure in our model of the dengue epidemic network, source of infection transfers from any other cluster (say *Cj*) onto *Ci.* For example, travelers originating from the countries that have dengue ailment may be an external factor to the uninfected areas as they are carriers of the disease. Besides the international travelers there may have an impact on locals who travel from one place to another. Especially, dengue infected people travel into the place where the dengue virus is absent. These people might therefore be a cause of the spread of dengue virus. Similarly, any simple mosquito that gets the dengue virus from a dengue infected person and passes it on to an uninfected person (Fig. 1) also acts as an external source (Fig.3). As described earlier, a dataset of Selangor has been utilized in this study and it is important to note that Kula Lumpur airport is used for international travelers here, and from the result analysis it has been noticed, besides the other places in Gombak, there are many dengue cases especially in the area of Batu Caves (a tourist spot) [13]. In Fig. 3, consider Gombak as cluster *Ci* that contained various infected and uninfected nodes. On the other side, cluster *Cj* is the global cluster having some infected (such as: *Lj1, Lj2*) and uninfected nodes (such as: *Li3, Lj6*). If any node in *Ci* gets infected through a node from cluster *Cj* (other than *Ci*), it is named as the influence of external factor and called case 2. For this, *Cj* infects *Ci* externally; for instance, *Cj* is Ampang and *Ci* is Gombak, where Ampang is a neighbouring place of Gombak.

**Fig. 3.**
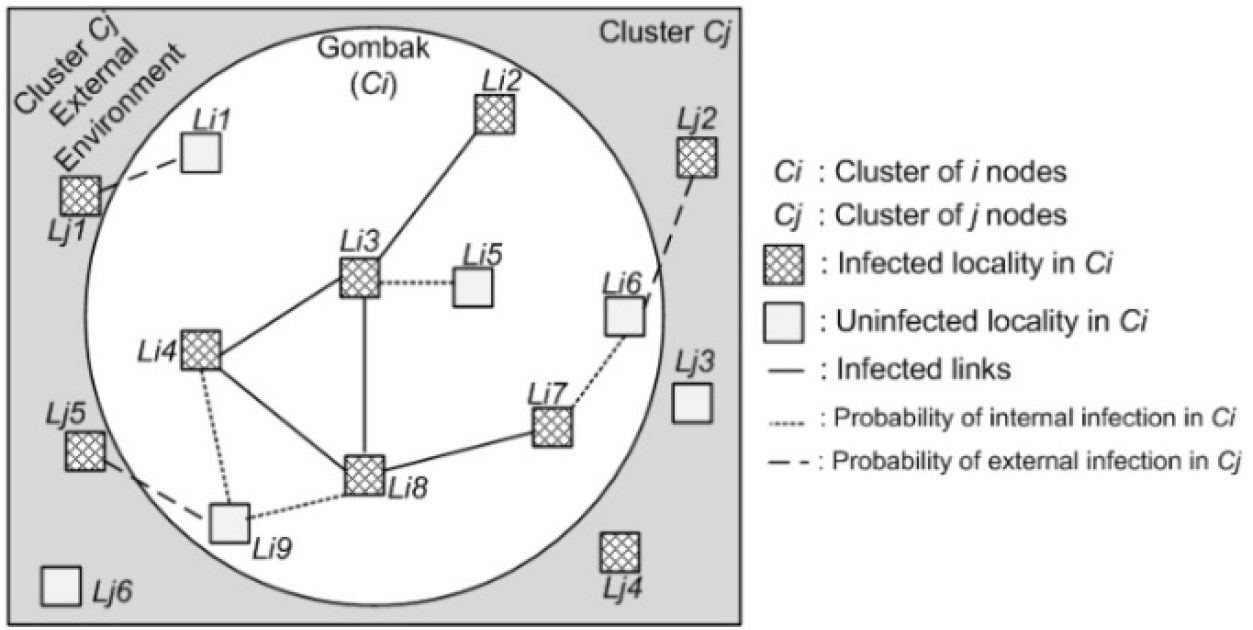
Influence of External diffusion (Scenario 1: Case 2), based on the given dataset.

### Scenario 2

What external factors can be inserted in the current dengue epidemic network to control the diffusion of *Aedes aegypti*? Sprays, cleanliness, vaccination and immunization all are the traditional methods and are being used regularly [19] [14] By measuring the prominent features of the scale-free network such as: Power-law, clustering coefficient, short path length, and by applying some centrality measures like, Degree, Betweenness, Closeness, Eigenvector, it is concluded that dataset of the dengue epidemic network has the topology as a scale-free network [12] [17]. That means there are few areas which work as focal hubs and by using different methods, these hubs are highlighted in the investigation and robustness has been observed [20]. Genetically Modified (GM) mosquito is an advanced technique which is centered on innovative genetic science [21] [22]. In this technique the DNA of the mosquito is changed to stop their generation growth. Here, only male mosquitoes (*Aedes aegypti*) are prepared by changing their DNA which mate with females later. Different countries have tried this new technique [23] [24]. Genetically Modified mosquito technique is preferred to be utilized in focal hubs as external factor to control the wild population of *Aedes aegypti* [25]. (Fig.4). The female *Aedes aegypti* mosquitoes that will have mated with GM male mosquito, lay eggs which later hatch into larvae; the offspring carries the piece of DNA which kills them before they become adults. The life span of the GM mosquito is up to 6 days, after this it dies [24]. The insertion of GM mosquito as an exogenous factor in the dengue epidemic network can produce efficient results especially when the topology of the network is scale-free nature. Furthermore, this way would be cost effective too. In Fig. 4, it has been shown that GM mosquito should be released in most infected nodes. It is referred to scenario 2 as ‘influence of external suppression’.

**Fig. 4.**
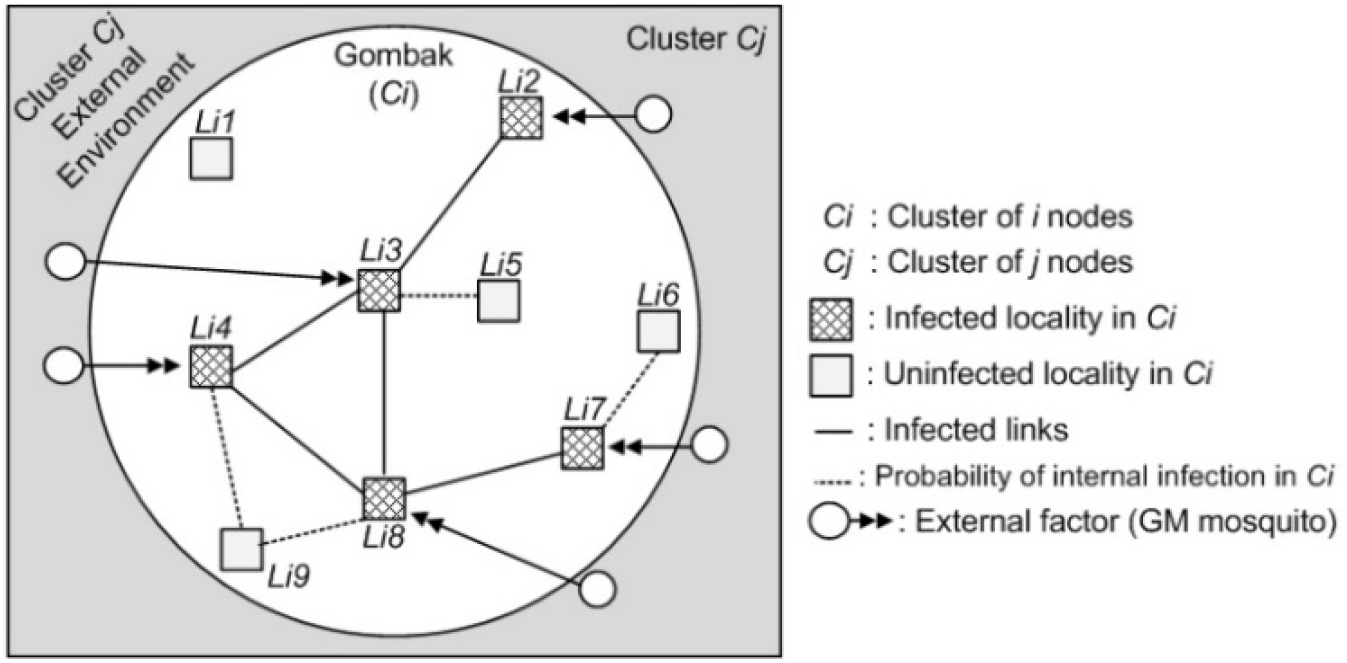
Influence of external Suppression (*Scenario 2*)

### 2.1 Modelling the infection influence

To model the emergence of dengue epidemic network in Selangor, there is a need to ponder on the movement of the undetectable exogenous source of this network that transforms the dengue virus to the nodes of the dengue network (via infected person or mosquito). The probabilistic generative model of virus occurrence is utilized in the dengue epidemic network, in which a dengue virus can approach a node through the links (infected person or mosquito) of this network or via the influence of the external source. Exposure and infections have different meanings in the model used in this study [26]. When any node gets exposed to dengue virus (DENV) that means an exposure event has occurred, and an infection event occurs when any patient appears in a node with DENV. Exposure to the virus leads to an infection. Through two ways, a node can get exposed to virus. Firstly, *Li1* a node might be exposed to or becomes aware of DENV if any of its neighbours in the dengue epidemic network of cluster *i* (*Ci*), pass on a DENV. Hence, any uninfected node of *Ci* which is exposed by any other infected node within the *Ci*, it is referred as an “internal exposure” (Fig.2), where *Ci* contains the localities *Li1, Li2… Li_N_.* Secondly, *Li1* might be exposed to DENV through any other cluster *j* (*Cj*) which may be the neighbour of *Ci.* This activity of the external source is named as “external exposure”. Hence, here any node of cluster *Ci* is exposed to the node of locality of any other cluster *Cj*, where *Cj* contains the localities *Lj1, Lj2… Lj_N_.* It is alluded to the quantity of exogenous exposures after a period as the ‘event profile’. So, to build up the association between exposures and infections, the concept of the exposure curve is characterized that maps time that node *Li_N_* has been exposed to DENV, into the possibility of *Li_N_* receiving infections.

By applying the virus diffusion model, we are able to consolidate the extent of virus from node to node along edges in the dengue epidemic network [17]. Event profile is proportional to the probability of any node (*Li_N_*) getting an exogenous exposure at a specific period [17] ‘contagion’ is used to mention a specific carrier evolving in the dengue epidemic network and when a DENV appears in an uninfected node it is referred as an infection of node by a contagion. These contagions are utilized jointly meaning these are considered together. The current model is delineated in a node-centric framework and it is assumed that after a time period, contagion becomes the reason to spread the intensity of exogenous exposure which a node gets and directed by the event profile *λ*_*ext*_(*t*). Moreover, by that contagion, the neighbours in the dengue epidemic network can also get infected which further become the cause of internal exposure. Every exposure has the probability to spread the infections to the uninfected node and the infection probability varies with the exposure curve *η*(*x*). Ultimately, it might generate any of the two possibilities that either exposure stops, or node gets infection and exposed to its next neighbour. It is also required to gather the amount of exposure which is produced by the exogenous means by time (*t*) also, in order to form the exposure curve *η*(*x*) which sees the probability of infection of node. Furthermore, we have given the modelling of internal and external exposures in *Appendix A.*

### 2.2 Data Analysis

For the modelling of internal and external exposure, we have utilized the data set given in *Appendix B*. That dataset has been divided into six clusters namely, Gombak, Klang, Hulu Langat, Petaling, Sepang and Hulu Selangor. The diffusion concept has already been explained in Fig.2 and Fig.3. Particularly, it can be observed that Hulu Selangor is in the neighbourhood of Gombak and Petaling. Therefore, cases appeared in Hulu Selangor (HS) are considered as internal exposure whereas with regard to HS, cases in Gombak are considered as external exposure and vice versa.

## 3. Results and discussion

Fig.5 presents the simulated results of above discussed scenario 1 (case 1 and case 2) which is related to the sections ‘modelling the internal exposure’ and ‘modelling the external exposure’. In this figure the dotted line shows the infections due to internal factors and the continuous line represents infections due to external factors, where x-axis shows the time in days. Here, the dengue infections are observed for 30 days, considering a month. On the y-axis the probability of infection is shown, which depends on the number of dengue cases that might appear in the *Ci.* Firstly, it is considered as case 1 (dotted line), the graph showed the maximum infections during 8 to 13 days. Possible DENV symptoms appear in 4 to 14 days. Here, it has been observed that the probability of infection approaching to maximize the value on the 11^th^ and 12^th^ day that is 0.8, meaning this time period is most crucial for the spread of DENV. This simulation has few conditions: 1-Time span is fixed up to 30 days. 2-It is supposed that there would be no external factors influence in case 1 network (like Fig.2). 3-It is assumed that *Aedes aegypti* bites only from day 1 to 5 (to see its maximum effects).

**Fig. 5.**
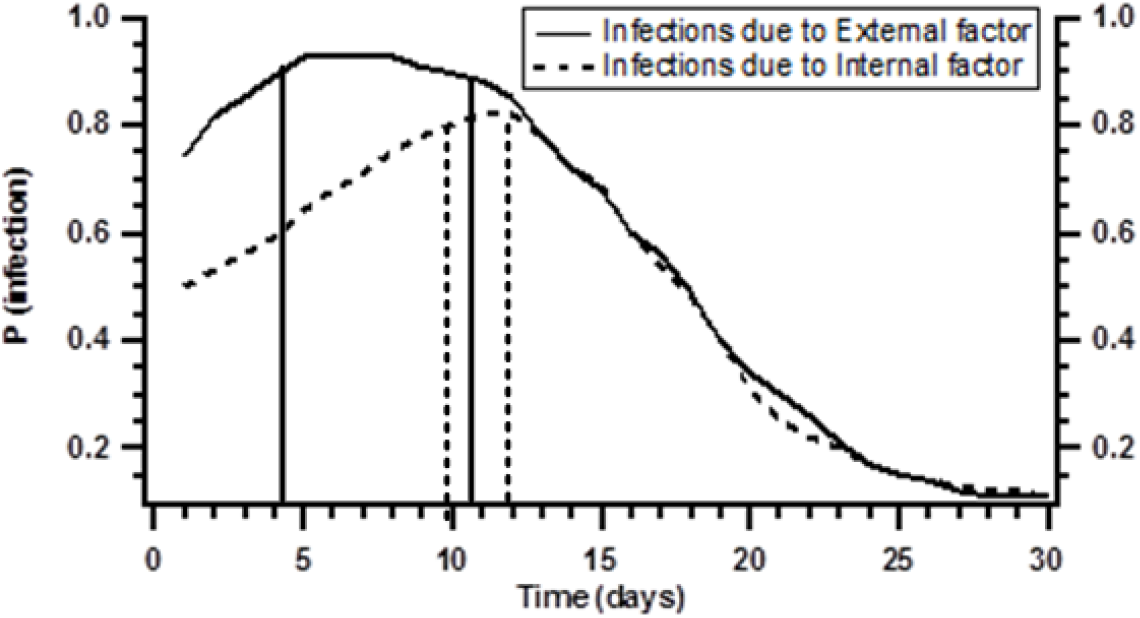
Comparison between case 1 and case 2 (*Scenario 1*)

Secondly, in case 2 (continuous line), the graph (Fig.5) clearly showed the increasing trend of the dengue epidemic network in the *Ci* as some external factors from the global cluster *Cj* influenced the network (see Fig.3). Here, the infection rate is high, and duration has been expanded due to the impact of external elements. In this case Infection range is high which is from day 4 to 14 and the peak of infection has been found during 5^th^ to 12^th^ day. Further, the probability of infection has reached between 0.9 to 1.0. This demonstrates how external factors have influenced the dengue epidemic network in Gombak Selangor.

Simplifying both cases (1 and 2) it can be observed that the peak of P(infection) = 0.8 in the first case whereas in the second case, due to the addition of some external factors peak of P(infection) has raised and ranged between 0.9 to 1. More specifically, P(infection) in the case of internal exposure is at the peak only, on two days: the 11^th^ and 12^th^ day. While in the external exposure case it is at its peak from 5th to 12th day and this indicates that infection time span has increased. In case 2, peak in the graph has been increased and shifted on the left side as well, that means external factors have expended the infection duration and maximized the probability of infections as well.

Fig. 6 is the simulated representation of the scenario 2 (shown in Fig. 4), where it was described how DENV might be suppressed by any external factors and how the chain network (Fig. 1) of dengue virus infection can be broken down. In the graph (Fig. 6), x-axis shows the time in number of days up to 34, as the average life span of *Aedes aegypti* is 34 days. While y-axis shows the probability of destruction of the dengue epidemic network. Here, the simulated results are shown by considering 1000 *Aedes aegypti* mosquitoes in the epidemic network. Five hundred Genetically Modified (GM) mosquitoes are inserted/ released on the network of *Aedes aegypti* (it has been discussed that only female *Aedes aegypti* can spread the DENV and GM mosquito are only male mosquitoes with changed DNA. The life span of GM mosquito is up to 6 days). The result showed that as 500 GM mosquitoes were released in the *Aedes aegypti* network, they mated with *Aedes aegypti* females (say 600 females) that means these 600 females were now unable to produce a new generation. These 600 females *Aedes aegypti* laid eggs, and larvae died before they grow up, because of the changed DNA characteristic of GM mosquitoes. The graph showed almost 60% network is destroyed in 5 to 6 days. Moreover, female *Aedes aegypti* mates with any male mosquito only once in her life [27]. Furthermore, after 6 days GM mosquito dies [27]. As a result of this, 60% of female mosquitoes cannot reproduce its new generation and 40% remaining female *Aedes aegypti* could live their average life up to 34 days before being finally wiped out from the network. Ultimately, infections remain very low and finally may be removed from the network and threshold value is achieved. There are some simulation conditions in this network: the numbers of days are fixed up to 34, number of *Aedes aegypti* females are 1000 in the fix time interval and the released 500 GM mosquitoes work properly.

**Fig. 6.**
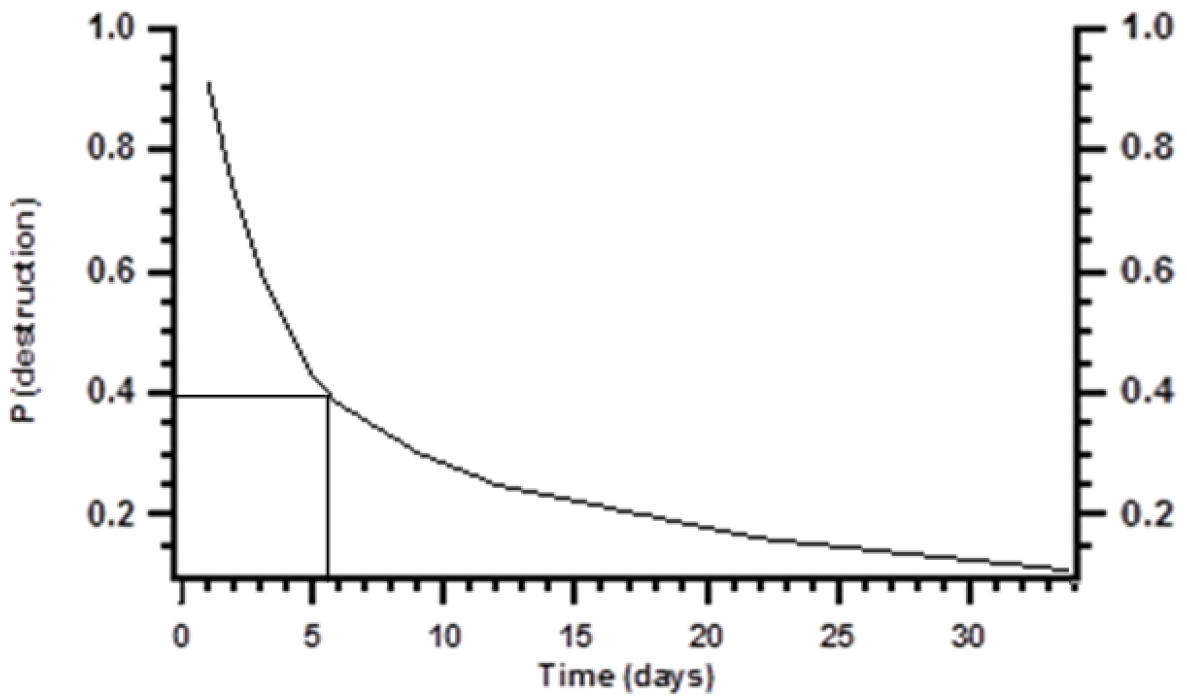
Destruction of network due to external factor (*Scenario 2*)

## 4. Conclusion

Despite of many regular methods such as aerosol, cleanliness and immunization *Aedes aegypti* is still uncontrolled, and is increasing exponentially, particularly in tropical zone in the world. Here, the complex network of dengue epidemic phenomenon has been analyzed through two aspects. Firstly, different factors that could be the reason of dengue virus diffusion are discussed. The spread can be due to internal factors or/and external factors (Fig.2 and Fig.3). Secondly, various elements are described which can control or suppress the wild population of *Aedes aegypti* (Fig.4). Here, the internal and external exposure of dengue epidemic network is modelled (*Appendix A*) and simulated results are presented for both factors. An increase in the dengue infections has been seen, as some external factors act in the *Aedes aegypti* network. External factors influence not only increased the probability of infection but have also expanded the infection intervals. Further, the results of the insertion of GM (genetically modified) mosquito into the *Aedes aegypti* network showed the destruction of the network (Fig.6). Hence, it is recommended that this technique will work more efficiently in the scale-free network through targeted attacks in the focal hubs of *Aedes aegypti*. Additionally, by considering the spatio-temporal technique, from the results of internal and external exposure modelling the best time period has been observed that could be utilized to suppress the dengue epidemic network. If a specific ratio of GM mosquitoes could be release at the particular time in specific nodes then that could produce better outcomes as compared to random procedure. *Aedes aegypti* is also the cause of *Zika* virus (ZIKV) which has become a serious threat for the whole world. Furthermore, this can be applied on ZIKV and other arboviruses to control these. Moreover, we have aim to apply this methodology on the CoViD-19 to see its network propagation and to observe the internal and external factors for its growing transmission.

## Appendix A

Modelling of internal exposures and external exposures

## Modelling the internal exposures

There is some important information regarding DENV that has been used in the model of the dengue epidemic network. Scientists found four serotypes of dengue virus namely, DENV-1, DENV-2, DENV-3 and DENV-4 [1] [16]. All these dengue serotypes can be detected from the infected person’s blood samples. Any person who is infected by any one of these serotypes will never be infected again by the same [8]. The average life of the *Aedes aegypti* is 34 days and when *Aedes aegypti bites* any person the symptoms appear in 4 to14 days [8] [12].

In this model, internal exposure is considered only when virus is exposed within the same cluster (say *Ci*) and then DENV infections appear. Exposure transmission has no fixed time limit, as nodes are exposed after random time interval *t*. Let’s observe an example from the real dengue epidemic network. Gombak is a cluster *Ci* of the said network that contained the various infected and uninfected nodes, where different contagions act to spread the DENV (Fig.2). These contagions can diffuse the virus only in the closed system (local network).They can only affect their neighbours so then, and only then, there is internal diffusion propagated along the edge. Moreover, an infected entity that exposes the DENV to its neighbours can spread infections only once, as stated above DENV infected person cannot be infected again by the same serotype.

Let *λ*_*int*_ be the internal hazard function, for any neighbouring nodes *i* and *j*.

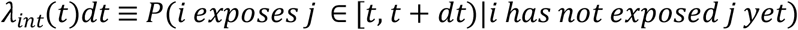

where *t* is the quantity of time that has gone since node *i* was infected. In our setting, *λ*_*int*_ efficiently models to what extent it takes a node to notice one of its neighbours getting infections [17] [21]. It is a component of the recurrence with which nodes observe each other. To Gombak dengue epidemic network, if any person is infected with dengue virus, the neighbour of that person is exposed by this virus. Means *Li*_*N*_ which is a neighbour of locality 2 (*Li2*) within the cluster *Ci* might be exposed. The probable amount of internal exposures, which any node *i* has received during time *t*, is defined as 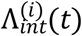, the total number of the cumulative distribution function of exposures propagating alongside with every node’s inbound edges and can be resultant as equation (1).

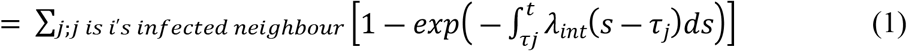

where *τ*_*j*_ is the time when node *j* gets infections.

## Modelling the external exposures

There may be many external factors that affect the whole dengue epidemic network. Here, some possible contagions are discussed that might be a cause of exposure in *Ci.* It has been a core issue that exogenous elements cannot be measured. External source does not have the same intensity. It changes from time to time; this function is referred to the ‘event profile’ which is defined as *λ*_*ext*_(*t*). Specifically, for any node *i*,

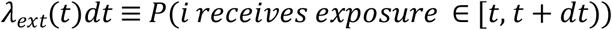

where *t* represents the quantity of time when the contagion appears for the first time in the dengue epidemic network [17]. A few points are to be noted. There is an equal probability for all nodes that they can get any external exposure at any time *t*. Secondly, there is a chance that any of the infected nodes may get the external exposure again. Simplifying this, any node may be infected more than once. As in the real dengue infection scenario, any infected person who gets DENV-1 serotype may be infected by DENV-2, DENV-3 or DENV-4. It is said *λ*_*ext*_ the event profile as it defines the real world event which caused the virus to reach the localities network and start diffusion. As the event grows over time *t*, its efficiency in the network varies.

Suppose that *λ*_*ext*_ is fixed for all time and that time is discretized into limited interims of length Δ*t*. At that point probability that *n* exogenous exposures have been occurred after *T* time interims is precisely a binomial distribution [17], such as equation (2).

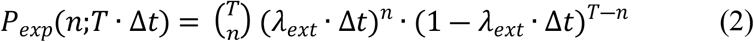

Set *t*= *T* ∙ Δ*t*. if the limit is considered as Δ*t*→0 and *T*→∞ such that *t* does not change. Then this probability approaches such as equation (3).

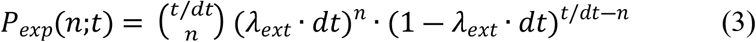

To relax the constraint that *λ*_*ext*_ is constant, average of *λ*_*ext*_(*t*) over *t* is used as equation (4).

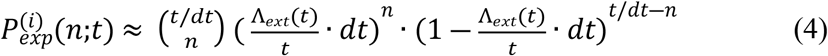

where 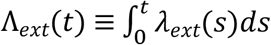. Eventually, nodes in the dengue epidemic network get both external and internal exposures at the same time, so both practices need to be taken into account. This would indicate receiving the convolution of the two probabilities that would not be feasible computationally. Instead, the mean value of 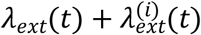 is used as equation (5).

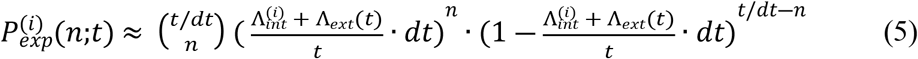

Productively, the unsteadiness of exposures is assessed as consistent in time such that every interim of time has an equivalent likelihood of an exposure arriving, so the entirety of the event is a standard binomial arbitrary variable.

## Appendix B

Dataset (20 October 2013 to18 October 2014) of dengue patients in Selangor, Malaysia. Dataset protects patients’ privacy.

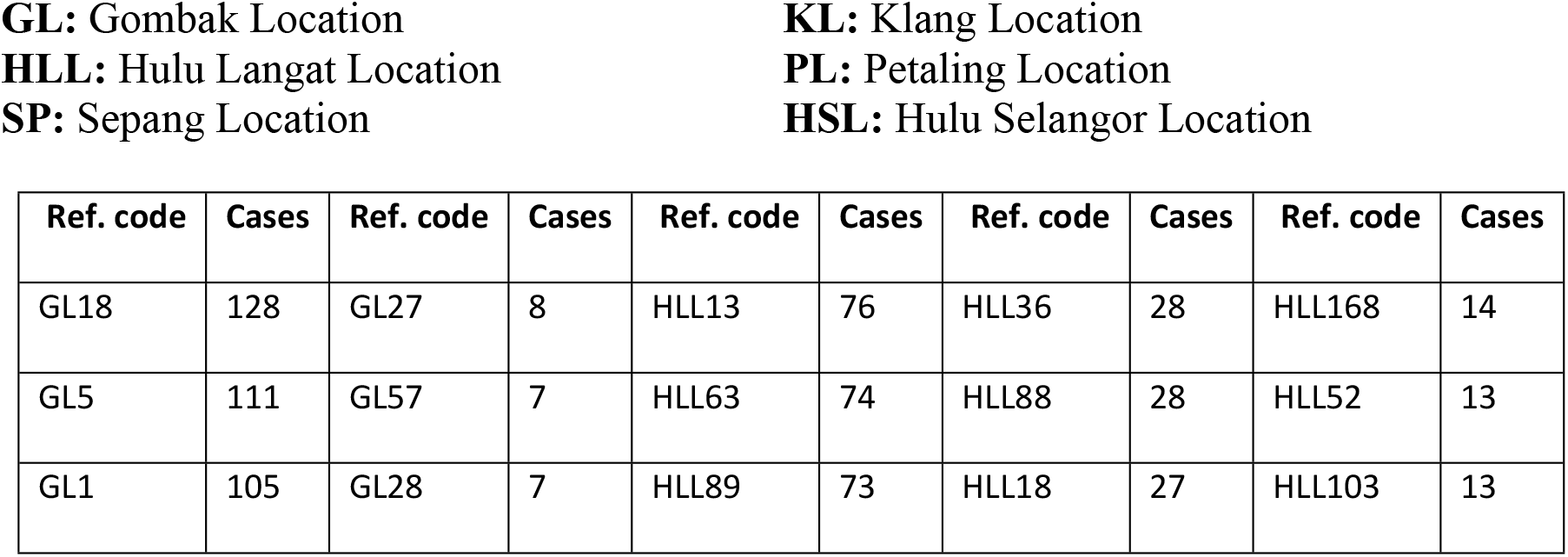
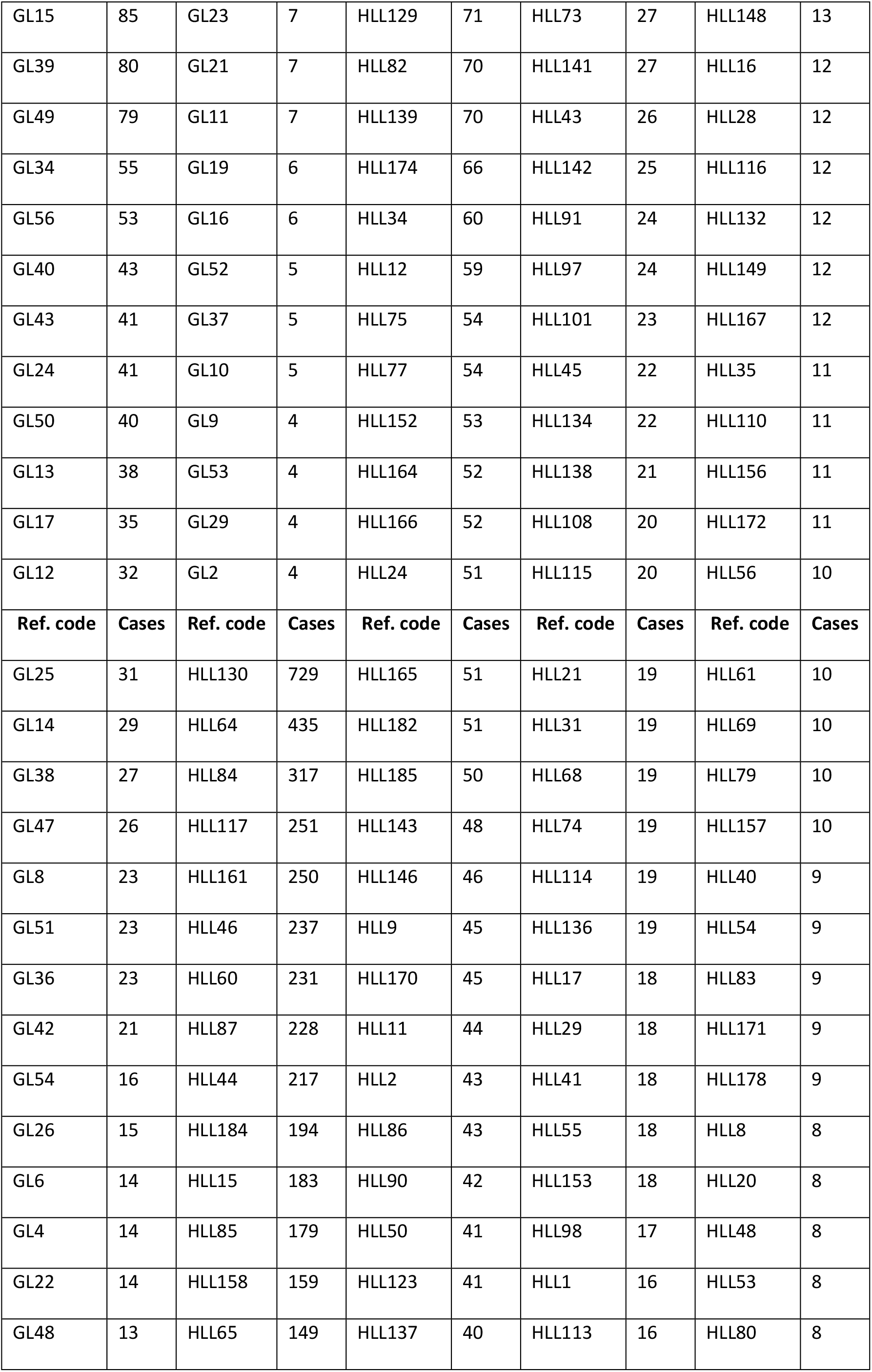
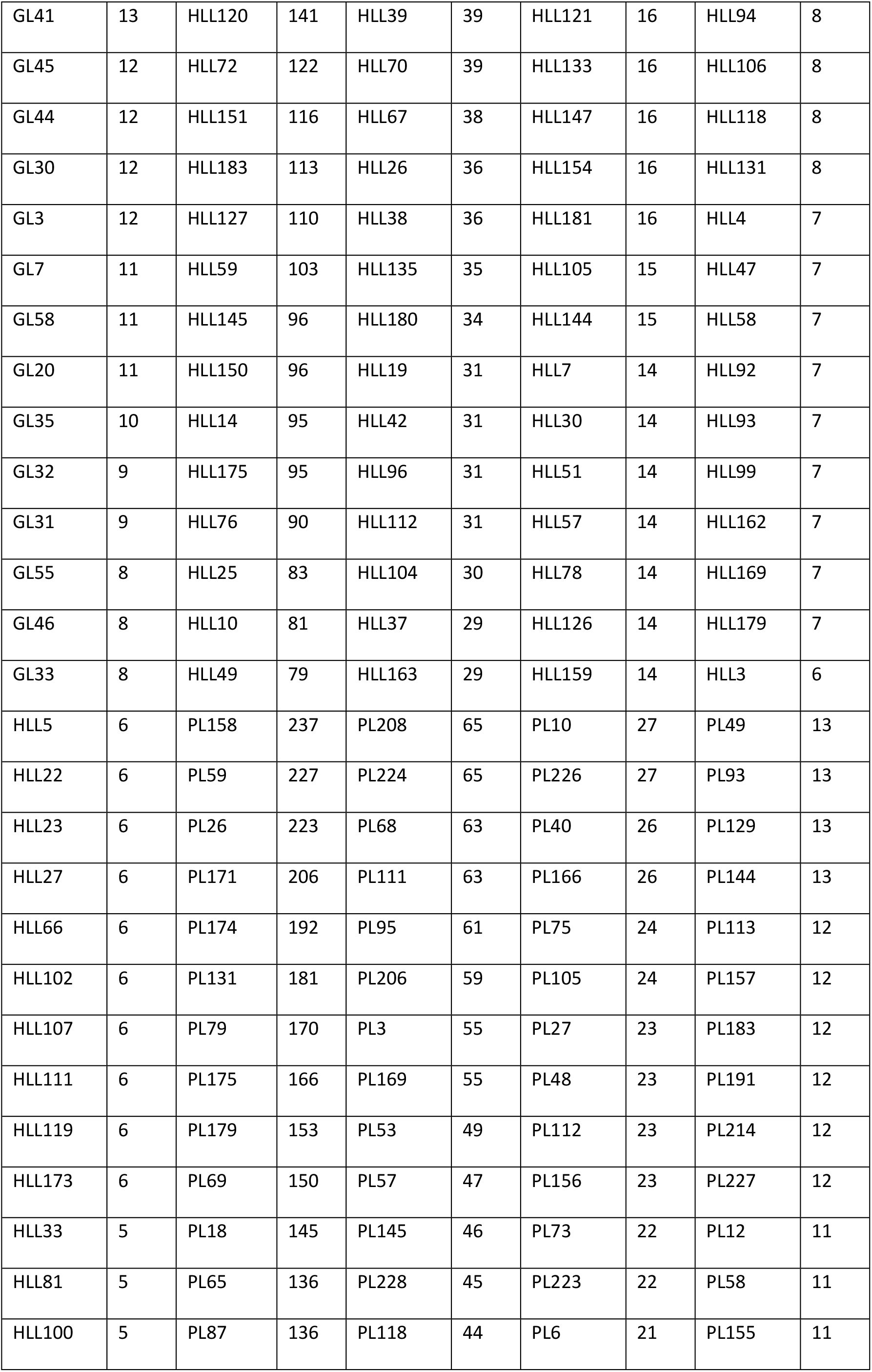
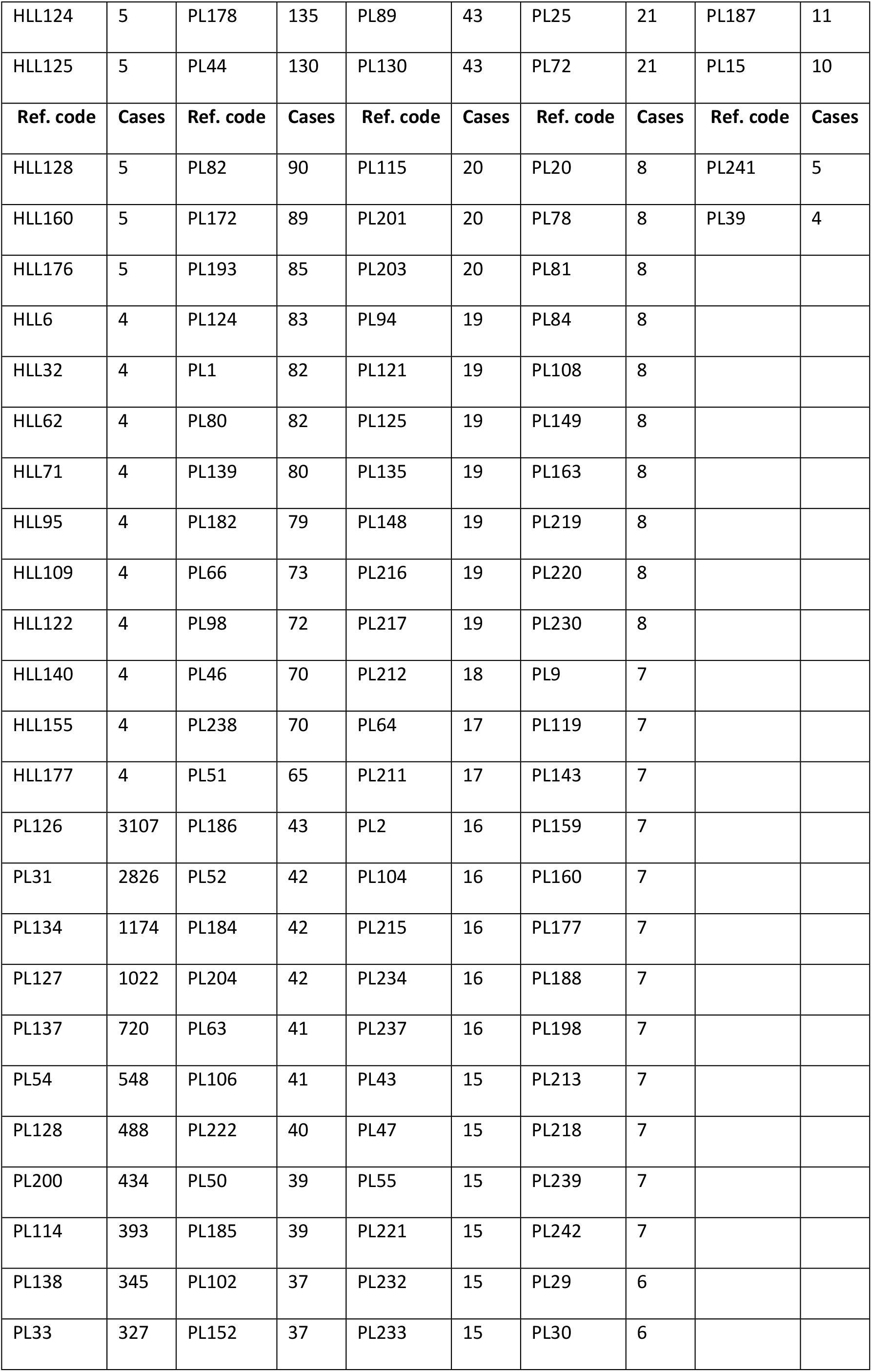
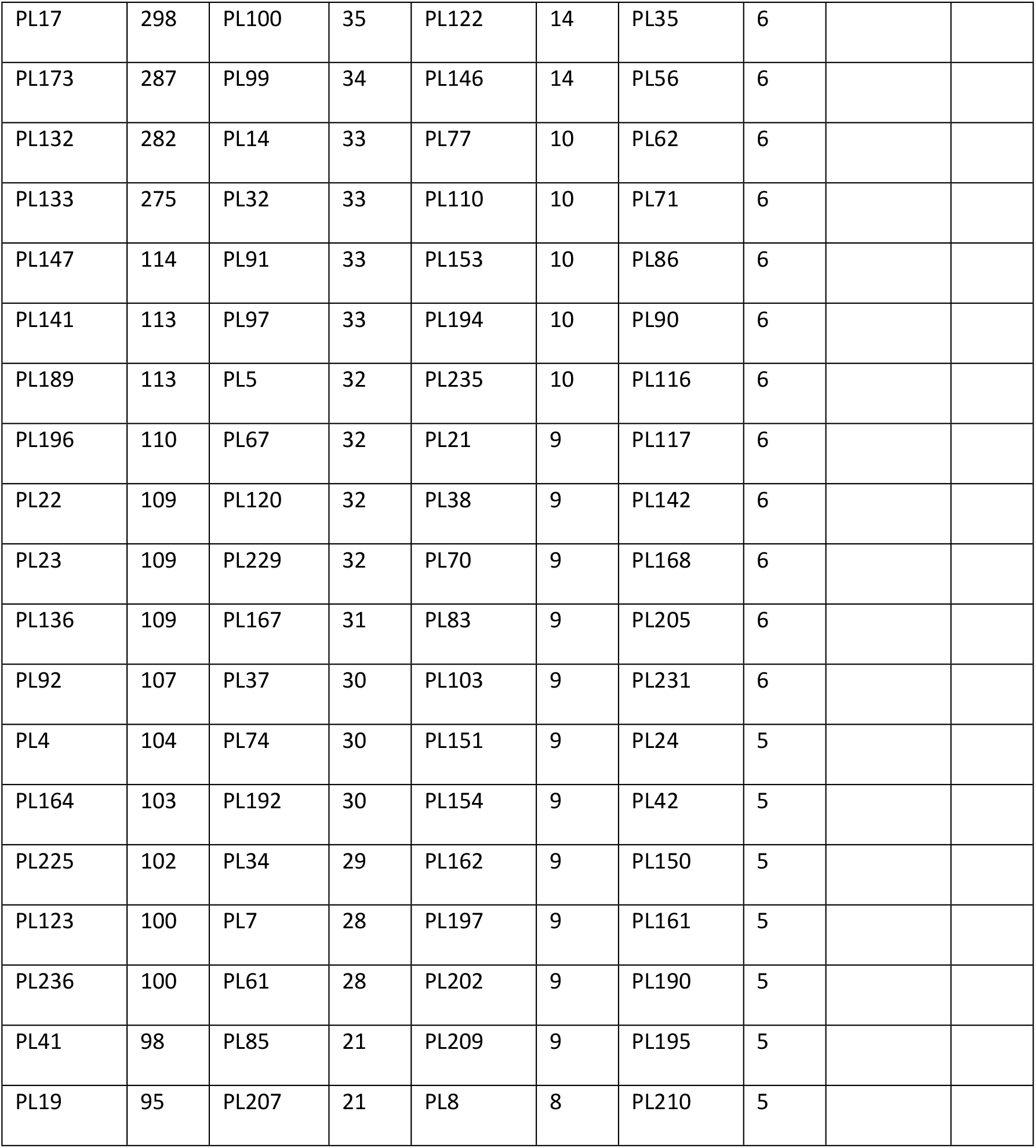

